# DISRUPTION OF THE ENDOGENOUS INDOLE GLUCOSINOLATE PATHWAY IMPACTS THE *ARABIDOPSIS THALIANA* ROOT EXUDATION PROFILE AND RHIZOBACTERIAL COMMUNITY

**DOI:** 10.1101/2023.11.29.569303

**Authors:** Daniel Acuña, Molly C Bletz, Joelle Sasse, Shirley A Micallef, Suzanne Kosina, Benjamin P Bowen, Trent R Northen, Adán Colón-Carmona

**Affiliations:** Department of Biology; University of Massachusetts Boston; Boston, MA, 02125, USA; Lawrence Berkeley National Laboratory, Environmental Genomics and Systems Biology, 1 Cyclotron Road, Berkeley, CA, 94720, USA; Joint Genome Institute, National Laboratory, Environmental Genomics and Systems Biology, 1 Cyclotron Road, Berkeley, CA, 94720, USA; Institute for Plant and Microbial Biology, University of Zurich, 8008 Zurich, Switzerland; Department of Plant Science and Landscape Architecture, University of Maryland, College Park, MD, 20742, USA; Center for Food Safety and Security Systems, University of Maryland, College Park, MD, 20742, USA

**Author notes:** Correspondence to: Adán Colón-Carmona; Department of Biology; University of Massachusetts Boston; 100 Morrissey Boulevard; Boston, MA, 02125, USA.

**Keywords:** Indole glucosinolates, root exudates, secondary metabolites, rhizosphere, root microbiome

## Abstract

Root exudates are composed of primary and secondary metabolites known to modulate the rhizosphere microbiota. Glucosinolates are defense compounds present in the Brassicaceae family capable of deterring pathogens, herbivores and biotic stressors in the phyllosphere. In addition, traces of glucosinolates and their hydrolyzed byproducts have been found in the soil, suggesting that these secondary metabolites could play a role in the modulation and establishment of the rhizosphere microbial community associated with this family. We used *Arabidopsis thaliana* mutant lines with disruptions in the indole glucosinolate pathway, liquid chromatography-tandem mass spectrometry (LC-MS/MS) and 16S rRNA amplicon sequencing to evaluate how disrupting this pathway affects the root exudate profile of *Arabidopsis thaliana*, and in turn, impacts the rhizosphere microbial community. Chemical analysis of the root exudates from the wild-type Columbia (Col-0), a mutant plant line overexpressing the MYB transcription factor *ATR1* (*atr1D)* which increases glucosinolate production, and the loss-of-function *cyp79B2cyp79B3* double mutant line with low levels of glucosinolates confirmed that alterations to the indole glucosinolate biosynthetic pathway shifts the root exudate profile of the plant. We observed changes in the relative abundance of exuded metabolites. Moreover, 16S rRNA amplicon sequencing results provided evidence that the rhizobacterial communities associated with the plant lines used were directly impacted in diversity and community composition. This work provides further information on the involvement of secondary metabolites and their role in modulating the rhizobacterial community. Root metabolites dictate the presence of different bacterial species, including plant growth-promoting rhizobacteria (PGPR). Our results suggest that genetic alterations in the indole glucosinolate pathway cause disruptions beyond the endogenous levels of the plant, significantly changing the abundance and presence of different metabolites in the root exudates of the plants as well as the microbial rhizosphere community.

## 1 INTRODUCTION

Glucosinolates are secondary metabolites present in most plants belonging to the Brassicaceae family, including several economically important crop cultivars such as broccoli, horseradish, kale, cabbage, rapeseed, mustard, rocket and canola as well as the biological model *Arabidopsis thaliana* (1–5). With more than 130 structurally different glucosinolates, these secondary metabolites are crucial for the plant’s fitness due to their important role in plant defense against insects, herbivores, biotic stressors and abiotic agents (4,6–12). In plant tissue, glucosinolates are compartmentalized and physically separated from myrosinase enzymes. When biotic agents disrupt the plant tissue, the myrosinase comes in contact with the glucosinolates, hydrolyzing them into active defense compounds, including isothiocyanates, thiocyanates, nitriles and epithionitriles (9,13,14). Interestingly, intact glucosinolates and their byproducts are exuded into the rhizosphere, where a shift in the abundance of microbes capable of using these metabolites as carbon sources has been observed (15–18). However, the exuded type and amount of glucosinolates is dependent on the plant species. In *Arabidopsis*, different natural accessions display genetic variations in the types and quantities of glucosinolates produced (9). Different natural accessions of *Arabidopsis* also produce different root exudate profiles that are associated with distinct rhizosphere bacterial communities (19). Root exudates contain primary and secondary metabolites. Primary metabolites include carbohydrates, amino acids and organic acids which bacteria can use as carbon sources. Secondary metabolites like flavonoids and auxins have also been shown to influence bacterial communities serving in various roles including as signaling molecules for the establishment of the rhizosphere microbial community, enrichment of beneficial microbes and as deterrent agents against pathogens (20–23).

Interestingly, some of the recruited PGPR induce systemic resistance through the jasmonic acid/ethylene pathway and salicylic acid pathways, resulting in the induction of aliphatic and indolic glucosinolates used for defense against herbivores and pathogens and as carbon sources for some microbes (24,25). Similarly, microbe-associated molecular patterns (MAMPs) trigger the production of tryptophan-derived indole glucosinolate metabolites such as camalexin, a phytoalexin that prevents pathogen infection (26). Camalexin has been shown to recruit the PGPR *Pseudomonas* sp. CH267 (27). Thus, there is an intricate relationship between glucosinolates produced and their role in the recruitment, establishment, and persistence of different microbes capable of using them as carbon sources. Therefore, we suspect that a possible disruption in the function of genes involved in the glucosinolate pathway could result in a shift in the glucosinolate profile of the plant, its root exudate profile, and the rhizosphere community composition.

In *A. thaliana*, the tryptophan-derived indole glucosinolates (IGs) are produced via a pathway that also intersects auxin biosynthesis. This pathway is positively regulated by the MYB transcription factor *ATR1* (28). This transcription factor regulates the genes *CYP79B2* and *CYP79B3*, functionally redundant cytochrome P450 enzymes involved in the conversion of tryptophan (Trp) to indole-3-acetaldoxime (IAOx), and *CYP83B1*, another cytochrome P450 enzyme that catalyzes the transformation of IAOx to *aci*-nitro compounds, which later become S-alkyl-thiohydroximates, the committed precursors of glucosinolates (29–32). An overexpressor line *ATR1* mutant, *atr1D*, upregulates these cytochrome P450 genes; conversely, the *cyp79B2cyp79B3* double mutant line has reduced production of IGs, camalexin and indole-3-carboxylic acid (ICA) (26,33,34). Herein, we evaluate the involvement of IG pathway alteration in the plant root exudate profile and in the establishment of the plant associated rhizobacterial communities using the overexpressor line *atr1D* and the *cyp79B2cyp79B3* double mutant. The *atr1D* line does not produce a conspicuous phenotype such as an elongated hypocotyl, bigger rosette or more adventitious roots when compared to the wild type (28). Conversely, the *cyp79B2cyp79B3* line produces a smaller hypocotyl, reduced rosette and reduced formation of adventitious roots (35). We hypothesized that we would detect changes in the root exudate profiles and significant difference in the relative abundance of distinct metabolites between the different lines. Moreover, we expected to observe a shift in the rhizosphere microbial community associated with plants containing different capacity for glucosinolates production.

## 2 MATERIALS AND METHODS

### 2.1 Experimental design

Loam soil was collected from a site at the Waltham UMass Field Station in Waltham, MA. The soil had not been exposed to pesticides or herbicides and was stored in the dark at 4°C. Potting soil used in this study was autoclaved, ensuring the suppression of any preexisting microbes in the soil and *A. thaliana* growth. The sterility of the autoclaved soil was verified by performing genomic DNA extraction as described below, and endpoint PCR with the same primers used in the study. The absence of amplicon bands in 2% agarose gel confirmed the effectiveness of the autoclaving process. Soil used in all experiments to grow the different plant lines was prepared by mixing 10 g of Waltham’s loam soil containing the native microbial inoculum with 35 g of previously autoclaved potting soil. This soil blend, referred to as bulk soil in this study, was prepared simultaneously for all the used plant lines by thorough mixing.

The glucosinolates mutant lines *atr1D* and *cyp79B2cyp79B3 double mutant* were provided by Dr. Judith Bender from Brown University. The wild-type Col-0 was obtained from the *Arabidopsis* Biological Resource Center (ABRC), Columbus, OH. Growth conditions were as described by Micallef et al., (19). Briefly, seeds were sterilized 33% v/v bleach for 10 min, washed and imbibed with sterile water at 4°C in the dark for 48 h. Seeds were plated in Petri dishes containing half-strength Murashige and Skoog (MS) medium, 0.9% w/v phytoagar and supplemented with 1.5% w/v sucrose, pH 5.7. Seedlings were grown in the plant room for two weeks at 22**°**C under short-day conditions (12 h of light at approximately 130 μmol photons m^−2^ s^−1^). A total of 24 seedlings from each of the plant lines were transplanted into soil. We used a block design approach, with four replicates of each plant line, with each replicate consisting of 6 seedlings (Figure S1).

### 2.2 Collection of Rhizosphere soil

Seedlings were transferred to pots containing bulk soil and covered for one week to ensure humidity. Seedlings were grown at 22°C under short-day conditions in the plant room and watered with sterile, deionized water. Four weeks after transferring seedlings to soil, plants were uprooted, and roots were shaken to remove excess soil. Seedling rhizosphere soil belonging to a genotype block was collected by scraping with sterile pipette tips onto sterile Petri dishes and stored in sterile Eppendorf tubes. All soil samples were stored at –20°C until rhizosphere DNA was extracted using a QIAGEN DNeasy PowerSoil Kit (Qiagen, Inc., Valencia, CA, USA) following the manufacturer’s protocol.

### 2.3 Exudate sample extraction for metabolomics analysis

Exudates from the three different lines were collected from a different set of plants than those used to study the rhizobacterial microbiome to ensure the absence of metabolites not produced by the plants. Seeds from the three different plant lines were sterilized in 33% v/v bleach for 10 min and thoroughly washed with sterile water. Seeds were imbibed in sterile water and place at 4°C in the dark for 48 h prior to plating them in rectangular Petri dishes containing 50 mL of sterile, half-strength MS medium containing 0.9% w/v phytoagar and supplemented with 1.5% w/v sucrose, pH 5.7. Petri dishes were placed in the plant room, and after 7 days of growth at 22°C under long-day conditions (16 h of light at approximately 130 μmol photons m^−2^ s^−1^). After 7 days, under a laminar flow hood, seedlings were moved into individual flasks containing sterile half-strength liquid MS medium supplemented with 1.5% w/v sucrose. For each plant line there were three biological replicates, each replicate contained 45 seedlings. Flasks were shaken at 170 rpm for 18 days at 22°C in long day conditions along with flasks containing only liquid media as a control. After 18 days, under a laminar flow hood, plant roots were thoroughly washed with sterile deionized water to remove all traces of liquid media and subsequently placed in flasks containing autoclaved deionized water. Flasks were then shaken at 170 rpm for 3 days at 22°C in long day conditions in order to collect metabolites from the root exudates. Flasks with only deionized water were used as blank controls and shaken for the same time period. Roots exudates and blanks were filtered using a 25 mm diameter syringe and 0.22 μm HPLC grade nylon filters (Corning Inc., Germany) to remove debris. Samples were then flash frozen in liquid nitrogen and lyophilized using a Labconco FreeZone lyophilizer (Figure S2). Lyophilized samples were resuspended in 1 ml LC–MS grade methanol (CAS 67-56-1; Honeywell Burdick & Jackson). Samples were sonicated for 15 min in a water bath at 23°C and centrifuged at 5,000 g for 5 min at 4°C and then filtered through a 0.22 µm filter. Supernatants were then transferred to new microcentrifuge tubes and using a speed vacuum evaporated at 25°C until dry. The root fresh weight was used to resuspend the samples at a 200 µl / 4.2 g ratio in LC–MS grade methanol with 15 μM internal standard (SI Table 1) (767964; Sigma-Aldrich).

### 2.4 Liquid chromatography–mass spectrometry methods and analysis

Metabolites were chromatographically separated using reverse phase chromatography on an Agilent ZORBAX RRHD Eclipse Plus C18, 95Å, 2.1 x 50 mm, 1.8 µm column. Eluted metabolites were detected using a Thermo Q Exactive Hybrid Quadrupole-Orbitrap Mass Spectrometer equipped with a HESI-II source probe. Chromatography and LC-MS/MS data acquisition was performed using the parameters specified in SI table 1. Targeted metabolite data were analyzed using Metabolite Atlas toolbox (https://github.com/biorack/metatlas) to obtain the extracted ion chromatograms and peak area (77,78). Targeted analysis was performed by comparison of detected mass to charge ratio, retention time and MS/MS spectra from samples to authentic reference standards (SI Table 2). Untargeted analysis was performed using MZmine2 (79) to extract features from sample files. The features list was filtered to remove isotopes and features without MSMS; from the filtered list, the most intense fragmentation spectra for each ion was uploaded to GNPS: Global Natural Products Social Molecular Networking (80) to obtain spectral matches from the online database. Features were filtered as follows: intensity threshold (ten times more intense than the extraction or media-blank controls), retention time (>1 min), cosine score (>0.7) and five or more matching ions. Additional filters were used to estimate average peak height (greater than one million counts) and two-fold difference between the treatment groups. Highly variable signals were filtered using the 5% lower bound of the peak height (calculated by the Student’s t distribution, greater than 1e^4^). Lastly, statistical significance was determined using pairwise-tests with p-value correction. All *p*-values were corrected using the Holm adjustment for multiple comparisons. Student’s upper and lower bounds were calculated using the Python Scipy Stats T module. Significance tests were performed using the Pengouin pairwise tests module, also in Python. The final features and their top hit (e.g., putative identification) can be found in SI Table 3. Statistical comparisons and boxplot for camalexin were prepared using R version 3.6.2, base packages and plyr1.8.5 (81), agricolae 1.3–5 (82) and Cairo_1.5-12.2 (83).

### 2.5 16S rRNA amplicon sequencing, processing and data analyses

Soil rhizosphere was grouped by blocks, for a total of four blocks per plant line, as previously described. After genomic DNA was isolated from soil samples, DNA concentrations were checked using a Thermo Scientific™ NanoDrop™ 2000/2000c Spectrophotometer. Aliquots from all samples were used to make working aliquots of 50 ng/µL DNA. PCR amplification and library preparation were carried out to set up the sequencing of the amplicons using an Illumina MiSeq 1×151 Reagent Kit v3 following the Earth Microbiome Project (EMP) protocol (http://www.earthmicrobiome.org/) (84). Briefly, the V4 region of the 16S rRNA bacterial gene was targeted using barcoded primers 515F (85) (5’-GTGYCAGCMGCCGCGGTAA-3’) and 806R (86) (5’-GGACTACNVGGGTWTCTAAT-3’) at a 10 µM concentration and amplified using ThermoFisher Platinum Hot Start master mix. PCR products have an expected size of 300-350 base pairs which were verified using 2% agarose gels. Amplicons were cleaned using Roche KAPA pure beads at 0.8X as suggested by the manufacturer. Amplicons were eluted in Tris buffer and quantified using Promega QuantiFluor® dsDNA. Pooled amplicons were sequenced on an Illumina MiSeq at the University of Massachusetts, Boston’s Center for Personalized Cancer Therapy Genomics Core.

### 2.6 Sequence processing

Demultiplexing, quality filtering, sequence clustering into sub-operational taxonomic units (sOTUs) and data processing was carried out using Quantitative Insights Into Microbial Ecology QIIME ^TM^2. -version 2018.11 (87). Briefly, sequences were demultiplexed and quality-filtered using default settings and clustering of sOTUs was performed using Deblur (88). Taxonomy assignment was performed using the naïve Bayesian classifier and RDP database (89,90). A phylogenetic tree was built using the fasttree feature from QIIME2^TM^ (91). The lowest sample had a sequencing depth of 40,000 reads; thus, all samples were rarefied to this depth. Alpha diversity metrics, sOTU richness and Faith’s phylogenetic diversity, as well as the beta diversity metrics, weighted and unweighted Unifrac were calculated in QIIME2^TM^.

### 2.7 Statistical analyses

Statistical analyses for the rhizosphere community dataset were performed using R version. 3.5.2 and the packages ecodist (92), qiimer2R (93), vegan (94), lsmeans (95), pairwise adonis (96), readr, dplyr, tidyverse and ggplot2 (97). The root microbial community alpha diversity – Faith’s phylogenetic diversity, sOTU richness and Shannon diversity – was analyzed using a general linear model (GLM), ANOVAs, and *post-hoc* pairwise comparisons. To investigate if community structure and composition (i.e., beta diversity) within each plant line was significantly different from one another, we calculated both weighted and unweighted UniFrac distances and ran permutation multivariate analysis of variance (PERMANOVA) models with the Adonis2 function in vegan. The weighted and unweighted UniFrac distances were visualized using principal coordinate analysis (PCoA). Homogeneity of the sample dispersion was checked with the Beta dispersion function in vegan. Linear decomposition model (LDM) (98) was used to better explain which bacterial taxa were enriched in the rhizosphere from the different plant types (alpha of 0.05 applied to the *q* value which is the adjusted *p*-value for multiple comparisons).

## 3 RESULTS

### 3.1 Shift on the root exudate profile of cyp79B2cyp79B3

To study how disrupting the endogenous glucosinolates levels alter the root exudate profile of the different plant lines, root exudates were collected and profiled using LC-MS/MS. We matched a total of 9000 features through untargeted analysis to putative IDs in the GNPS database in the three plant lines. After removing duplicates and matching them to a putative ID, we were left with 18 compounds significantly different between *cyp79B2cyp79B3* and Col-0, whereas we observed only one significant feature *atr1D* and Col-0 were compared.

Consequently, this comparison was removed from further analysis and discussion. (Figure 1). Specifically, there was a reduction of metabolites like the organic acids mandelic acid and sinapoyl malate, the nitrile 9-(Methylsulfinyl) nonanenitrile and the nucleoside 5’-Methylthioadenosine in the exudate profile of *cyp79B2cyp79B3* when compared to Col-0. There was also a reduction of malic acid, and due to their similar fragmentation spectra, it was labeled as malioxamycin, an antibiotic only produced by *Streptomyces lydicus* (36) and by *Streptomyces griseus* (37). We also observed an increase in metabolites like ICA, pimelic acid, and 8-hydroxyquinoline-2-carbaldehyde when compared to Col-0 (Figure 1). Additionally, we evaluated the relative abundance of camalexin in the root exudate of the three lines. We observed that the relative abundance of camalexin in *cyp79B2cyp79B*3 was significantly lower (*p*-value <0.05) than the levels in Col-0, whereas no significant differences were observed between *atr1D* and Col-0 (Figure 2).

**Figure 1.**
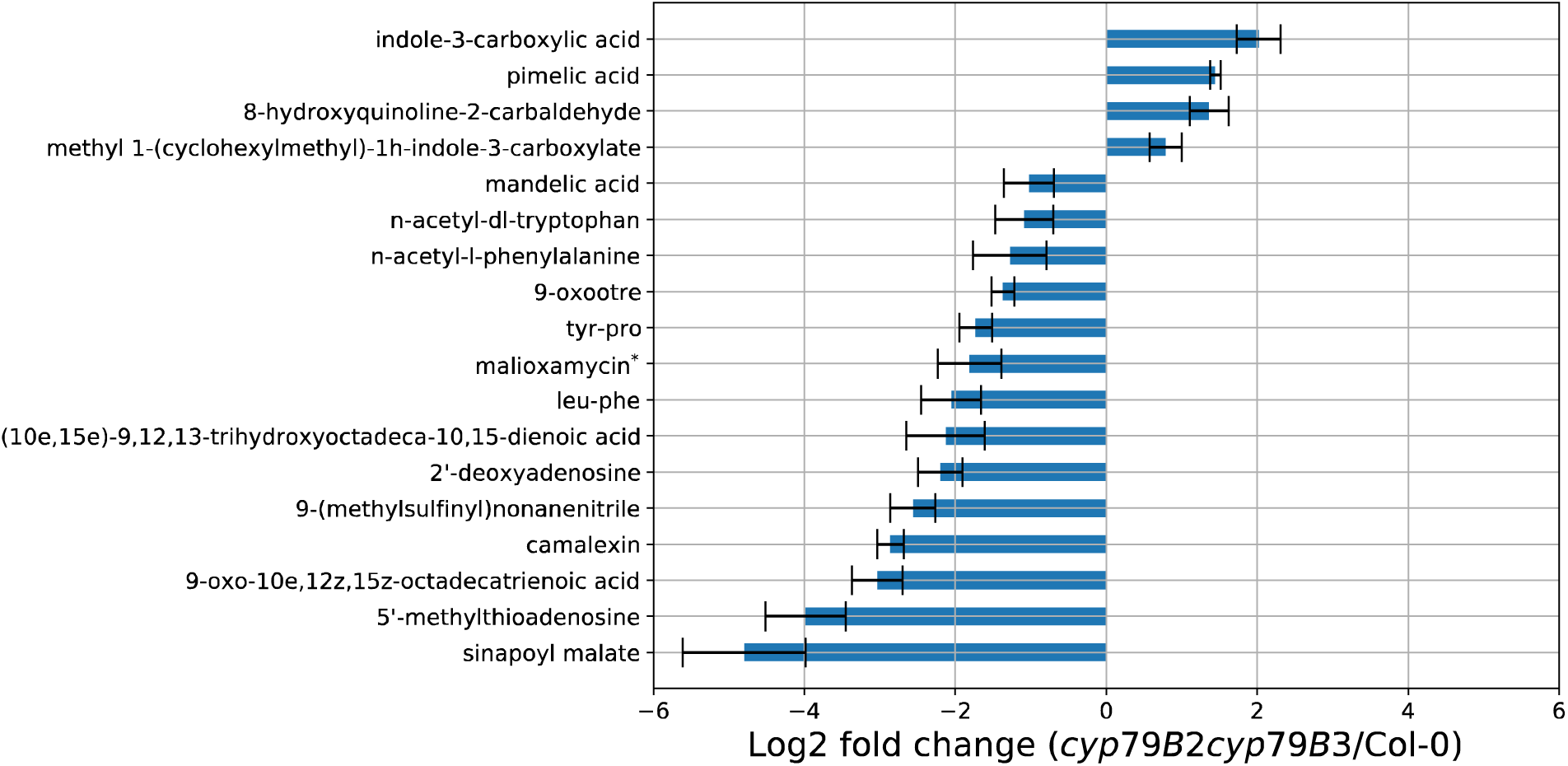
Putatively identified features with differential abundance between *cyp79B2cyp792B3* and Col-0 exudates (* the second-best match to reference spectra for malioxamycin is malic acid).

**Figure 2.**
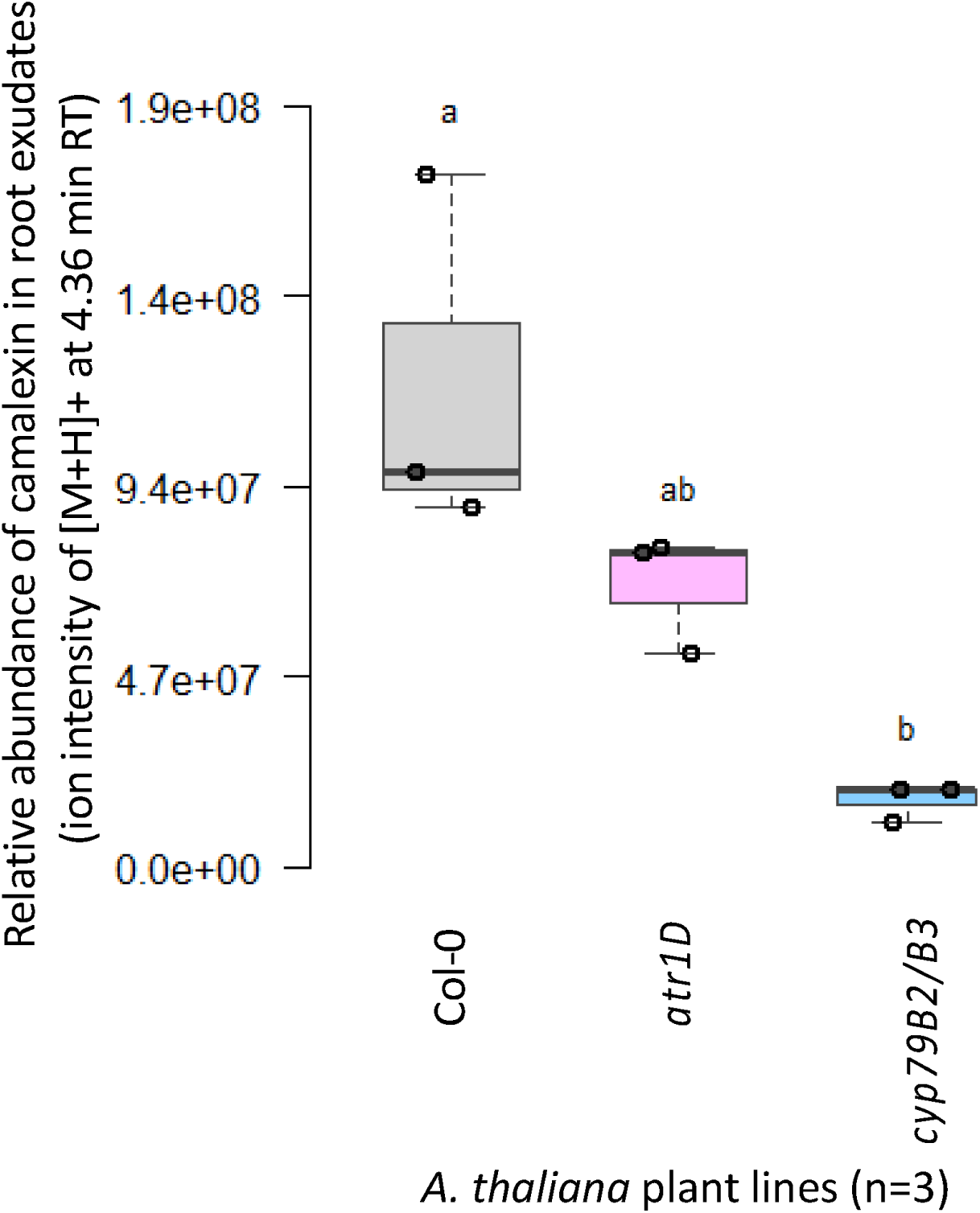
Relative abundance of camalexin in the root exudates of Col-0, *atr1D* and *cyp79B2cyp79B3* (*cyp79B2B3*). No significant differences were observed between Col-0 or between *atr1D* and *cyp79B2cyp79B3*. Camalexin levels were significantly lower in *cyp79B2cyp79B3* when compared to Col-0 (*p*-value < 0.05).

### 3.2 Rhizobacterial community diversity and composition

The most abundant phyla in the rhizosphere of the three *Arabidopsis* lines were Proteobacteria (51.90%), Bacteriodetes (10.20%), Acidobacteria (9.70%), Actinobacteria (8%), Verrucomicrobia (5.8%), Planctomycetes (5.15%), Gemmatimonadetes (3.90%), Chloroflexi (1.23%), Firmicutes (0.80%) and Armatimonadetes (0.70%). The relative abundance at the bacterial phyla level in the rhizosphere of the Col-0, *atr1D* and *cyp79B2cyp79B3* were similar. However, using LDM, we identified 24 specific sOTUs that were significantly different among the different lines. We detected higher relative abundance in sOTUs from the taxa *Oxalobacteraceae, Pseudomonas, Methylibium* and *Arthrobacter* in Col-0. sOTUs from the taxa *Acidobacteria, Myxococcales, Cyanobacteria, and Pirellulaceae* were higher in the rhizosphere of *atr1D* and sOTUs from the taxa *Flavobacterium* and *Solirubrobacterale*, *Devosia*, *Sinobacteraceae* and *Comamonadaceae* were higher in *cyp79B2cyp79B3* (Figure 3). Finally, we observed that the relative abundance of the taxa *Solirubrobacterales, Myxococcales*, *Opitutaceae*, *Deltaproteobacteria* and *Cyanobacteria* was higher in the mutant lines compared to Col-0.

**Figure 3.**
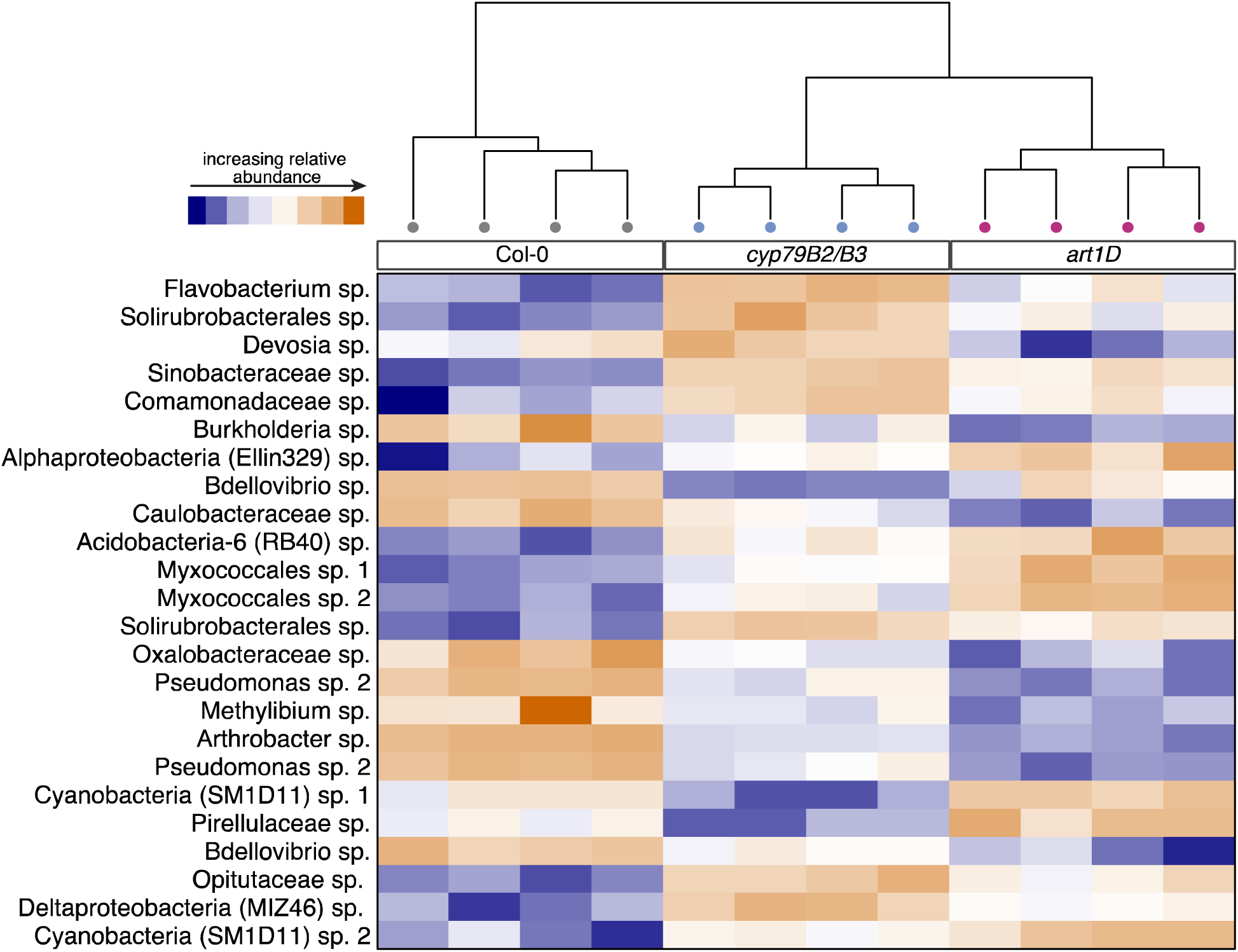
Linear decomposition model for differentially abundant sOTUs in the rhizosphere of Col-0, *cyp79B2cyp79B3* (*cyp79B2B3*) and *atr1D*. Twenty-four sOTUs were identified as significant (alpha = 0.05 applied to the q value which is the adjusted *p*-value for multiple comparisons).

We looked at two different alpha diversity metrics to determine the rhizobacterial richness and diversity within each of the plant lines. Using Shannon diversity analysis, we found that *atr1D* had significantly higher diversity than Col-0 and *cyp79B2cyp79B3* (LR Chi-square = 22.90, df = 2, *p*-value < 0.001) (Figure 4), whereas there were no significant differences between Col-0 and *cyp79B2cyp79B3*. Faith’s phylogenetic diversity was not significantly different (Figure S3). The community structure and composition also differed between plant lines. This can be seen in the clustering of samples from each plant line for both unweighted (Pseudo-F_2,9_ = 1.529, R^2^ 0.253, *p*-value < 0.001) (Figure 5A) and weighted Unifrac (Pseudo-F_2,9_ = 8.827, R^2^ 0.662, *p*-value < 0.001) (Figure 5B).

**Figure 4.**
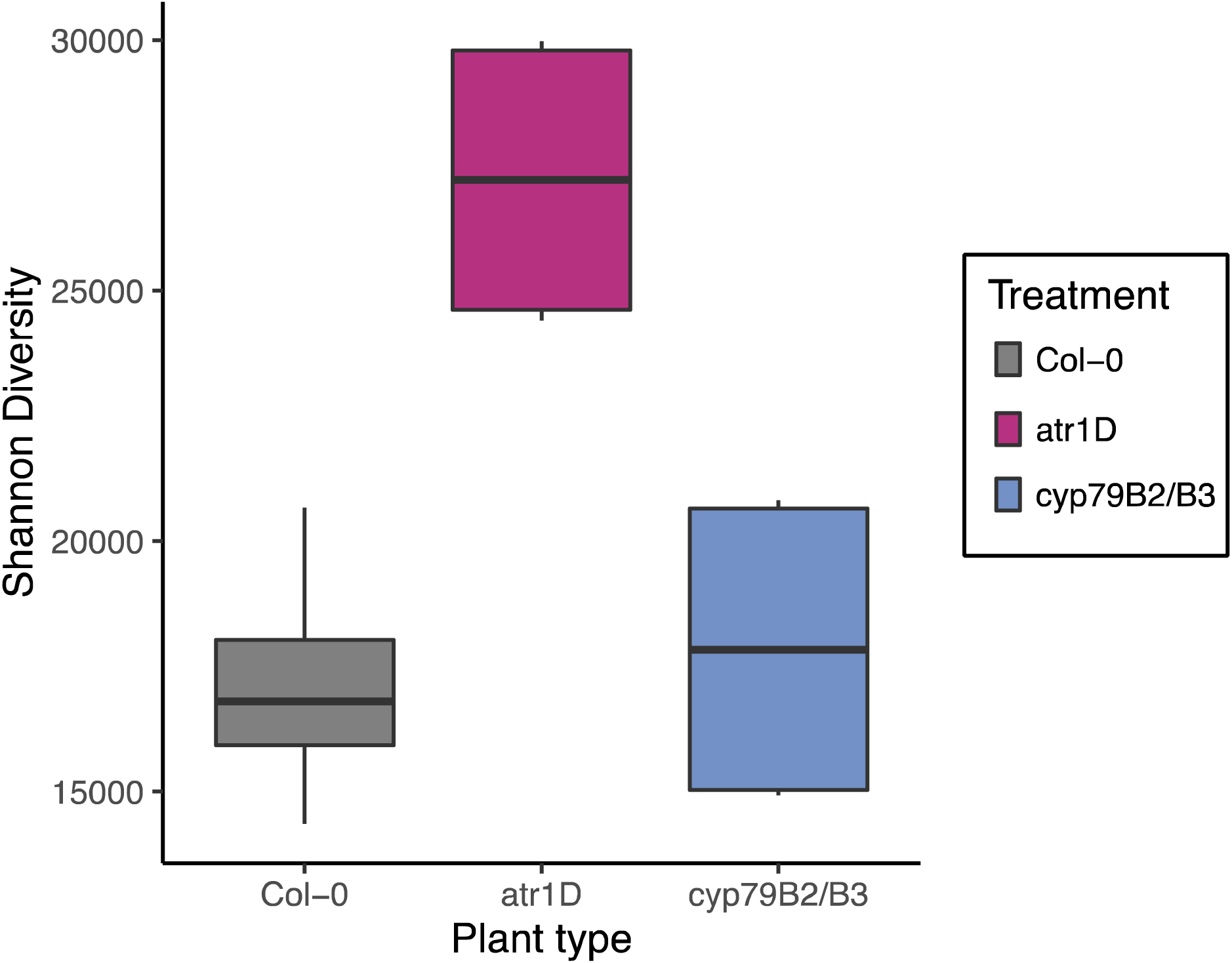
Differences in diversity between rhizobacterial communities from the different plant lines. The *atr1D* sOTU richness is significantly different from Col-0 and *cyp79B2cyp79B3* (*cyp79B2/B3*) *(p*-value < 0.0001).

**Figure 5.**
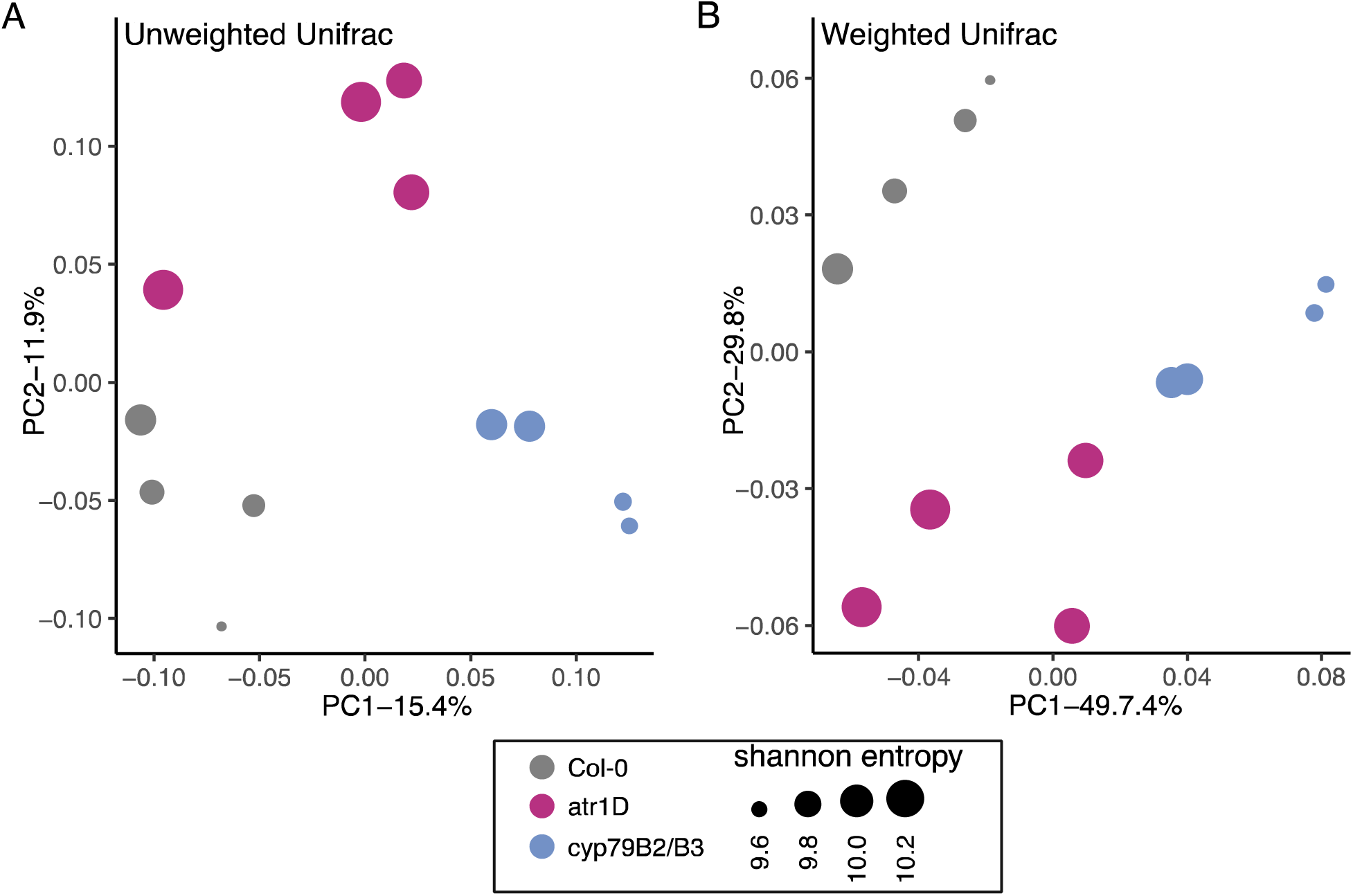
Principal coordinate analysis (PCoA) of rhizobacterial community composition from the different plant lines using (A) unweighted and (B) weighted UniFrac. Adonis results from the weighted and unweighted UniFrac analysis showed a significant difference between the rhizobacterial community from the different plant lines *(p*-value < 0.001).

## 4 DISCUSSION

Root exudate profiles are determined by plant genotype, developmental stage, and nutritional status. However, there are biotic and abiotic factors and pleiotropic effects caused by gene disruption that can alter the root exudate profiles (23,38). The combination of these factors is responsible for the variation in root exudate profiles in plants (39); and it is clear that root exudates are critical for the establishment of the rhizosphere microbiota (40–43). Rhizosphere microorganisms use some of the exuded metabolites as carbon sources. In turn, the associated rhizosphere microbiota provides functional benefits for the host, such as nutrient acquisition, stress tolerance and pathogen suppression (44). Due to this interconnected relationship, the rhizosphere microbiota will fluctuate in parallel with the developmental and nutritional statuses of the plant (45). Therefore, shifting root exudate profiles can directly affect the rhizosphere microbiota, a crucial player in plant health, pathogen defense and overall plant fitness (44,46,47). In our study, we used plant lines with altered indole glucosinolate pathways to investigate the role these secondary metabolites have in the establishment of the root exudate profile and rhizosphere community in *Arabidopsis*. We first collected and analyzed root exudates from Col-0, *atr1D* and *cyp79B2cyp79B3* grown hydroponically in axenic condition. Results from the overexpressing line *atr1D* suggest that upregulating glucosinolate levels does not cause a shift in the root exudate profile, at least features with putative IDs assigned. In contrast, the root exudate profile of *cyp79B2cyp79B3* was significantly different from Col-0 and *atr1D*. These results provided support for our initial hypothesis regarding the involvement of these compounds in establishing the root exudate profile of the plant and that there would be a shift in the profile when the indole glucosinolate pathway is altered. Analysis of camalexin were in agreement with results from Celenza et al., (28) where they also suggested that the synthesis and regulation for camalexin is not limited by IAOx and likely to be induced by pathogens.

Still, these exudate analyses may not fully represent the root exudate profiles of plants growing in the wild. Recruitment and interaction between the plant and microorganisms, especially pathogens and PGPR, elicit different metabolic pathways in the plant that ultimately affects the root exudate profile. Moreover, hydroponic conditions can alter the plant root morphology and architecture, which may result in changes in the root exudate profile (48). Thus, the actual root exudate profile from plant lines when grown in soil, in the presence of microbes, might differ from our findings. Nonetheless, our results help to better understand the differences and similarities we observed in the rhizobacterial community of the plant lines with different glucosinolate levels.

The three plant lines used in this study displayed higher relative abundance of specific bacterial taxa present in their rhizosphere. Col-0 had a high relative abundance of beneficial plant microbes such as Arthrobacter, known to produce indole-3-acetic-acid (IAA) and siderophores that influence root development (49), and bacteria from the *Oxalobacteraceae* family which are involved in nitrogen acquisition for the plant (50). We also observed two different species of *Pseudomonas.* Since *Pseudomonas* are one of the most predominant PGPR genera, we suspect that these species could be involved in the induction of induced systemic resistance to suppress foliar and root pathogens as well as to increase the presence of beneficial microbes in the rhizosphere of Col-0 (51–54). We hypothesize that these microbes are part of the plant’s core microbiome at the developmental stage at which we screened the rhizosphere of Col-0, given their properties and relative abundance.

The mutant *atr1D* displayed a significantly higher relative abundance of other potentially beneficial bacterial taxa. Among them were Acidobacteria, capable of producing IAA, siderophores facilitating nitrogen absorption (55) and Cyanobacteria, known to produce plant-growth hormones and fend off pathogens (56). In contrast, we also observed a high relative abundance of Myxococcales, known to be the most abundant and ubiquitous predatory bacteria in soil and capable of modulating the soil bacterial community (57). Hull *et al.* (31) had previously reported a 10-fold increase of IG levels in *atr1D* when compared to Col-0. Although the root exudate profile of *atr1D* was similar to Col-0 in axenic culture, we hypothesize that when *atr1D* is exposed to MAMPs in the soil, the root exudate profile shifts. This could be investigated in future studies where *atr1D* plants are grown in the same liquid media conditions as this study and treated with and without MAMPs to determine if the root exudate profile varies depending on the presence of MAMPs.

Another possible reason for significant differences in the high relative abundance of bacterial taxa in the absence of a root exudate shift between *atr1D* and Col-0 may be the constitutive up-regulation of key glucosinolate P450 genes in the *atr1D* mutant likely increasing the persistence of IGs in the rhizosphere, which may result in the higher abundance of taxa capable of using IGs as a carbon source. This last statement is supported by the significantly high alpha diversity observed in the rhizosphere of *atr1D* when compared to the other two lines (Figure 4). While it was beyond the scope of this study, future studies should focus on the levels of different relevant IGs such as indol-3-methylglucosinolate, 3-indoleacetonitrile, 2-phenylethyl isothiocyanate and 3-phenylpropanenitrile to better understand the complexity of the response.

We observed that *cyp79B2cyp79B3* also had a significantly higher relative abundance of specific beneficial bacterial taxa. Among them were *Flavobacterium*, which is used as biofertilizer (58)*, Devosia*, which is capable of producing IAA and siderophores (59) and *Comamonadaceae,* known to inhibit pathogen growth (60). We hypothesize that the high abundance of these bacteria is attributed to the overall resulting cocktail of exudates resulting from the disruption of the indole glucosinolate pathway. Nonetheless, we propose a correlation between certain exudates and the presence of different bacteria. The reduction of compounds that contribute to a stringent rhizosphere environment such as 9-(Methylsulfinyl) nonanenitrile (61) or mandelic acid (62) or 5’-Methylthioadenosine (63) could be compensated by the high presence of the antimicrobial quinoline 8-Hydroxyquinoline-2-carbaldehyde (64). Moreover, we observed a high abundance of pimelic acid, a biotin precursor that can potentially attract some, but not all, of the beneficial bacteria we observed (65), potentially having a similar effect to malic acid, a compound known to recruit beneficial bacteria (40). Future work could use this plant line to investigate whether the aforementioned bacterial taxa are recruited and confer benefits to the plant when challenged by different biotic and abiotic stressors. Interestingly, we observed a higher production of ICA in the *cyp79B2cyp79B3* mutant when compared to Col-0, which is puzzling. The monooxygenases CYP79B2 and CYP79B3 are necessary for the biosynthesis of ICA and its derivatives and is stimulated when the plant is challenged by biotic and abiotic factors (66). However, *cyp79B2cyp79B3* has reduced production of IGs, camalexin and ICA in the presence of pathogens (66). Given the vast battery of enzymes that plants have, this could suggest the existence of a Trp-independent pathway to produce ICA. For instance, there are different pathways used to produce IAA that do not rely on the monooxygenases CYP79B2 and CYP79B3. Such is the case of Trp-dependent pathways indole-3-pyruvic acid (IPyA) (67) and indoleacetamide pathway (68) to produce IAA. Conversely, a Trp-independent pathway that relies on the cytosol-localized indole synthase (INS) has also been reported to produce IAA (69). Moreover, the nitrile-specifier protein NSP hydrolyzes glucosinolates in *Arabidopsis* to produce nitriles (18,70), which seems to be advantageous to the plants when challenged by a biotic factor over the production of isothiocyanates (71). Consequently, we wonder whether the osmotic stress caused by the deionized water to collect root exudates could have triggered a mechanism similar to an herbivore attack. We speculate that, like IAA, there could be other pathways to produce ICA, whether as a byproduct of Trp-dependent IAA pathways or via NSP by converting glucosinolates to nitriles and other byproducts.

Plant root exudate profile change throughout the plant’s lifespan; along with changes, the rhizosphere microbial community that is associated with the plant (72). Rhizoengineering could modulate the rhizosphere community by means of enhancing or disrupting a metabolic pathway. Beta diversity results from this study confirm that disrupting the IGs pathway caused a clear shift in the rhizosphere community of the mutant lines (Figure 5). Targeting metabolic pathways in plants could be an interesting way to prime the soil with beneficial microbes as a step towards more sustainable agricultural practices. One example is the high relative abundance of bacteria from the taxa *Sinobacteraceae* in the rhizosphere of *cyp79B2cyp79B3*, capable of metabolizing herbicides present in soil (73).

The duality between the recruitment of both beneficial and potentially detrimental microbes into the plant’s rhizosphere highlights the influence that secondary metabolites have on the establishment of the rhizosphere community. These results also bring attention to the practice of using plant tissue from the *Brassicales* order as biofumigants to control soil-borne pests (16,74). Although environmentally friendly in theory, this practice inadvertently impacts beneficial soil microbes, involved in the nitrogen cycle as well as symbiotic microbes such as the arbuscular mycorrhizae (75). Interestingly, higher concentrations of root IGs may make the plant more susceptible to diseases such as clubroot infection, most likely due to the conversion of IGs to IAA, which is necessary for the gal formation (7). Plants resistant to this infection typically display a lower amount of root glucosinolates (7,76). Therefore, increased glucosinolate persistence in the soil could inadvertently increase the susceptibility of plants to different pathogens. The glucosinolate persistence and influence on soil communities’ idea is an interesting future direction for this work. We might hypothesize that multiple generations of plants growing in the same soils might have a similar pattern in the soil to what was observed for *atr1D*; an initial spike in species richness followed by a reduction of microbial richness due to bacterial communities being harmed by IGs. In sum, follow-up MAMP and multi-generational studies could provide a better understanding of plant-rhizosphere feedback systems and contribute to the fine tuning of agricultural practices when selecting break crops, as well as fostering and maintaining high microbial richness in soils to grow Brassicaceae crops.

## 5 CONCLUSION

The root exudate profile of the plants is determined by different factors such as the plant developmental stage, nutritional status or genotype. The exudate profile can be shifted by biotic and abiotic factors as well as gene disruption that in turn can directly affect metabolic pathways. Bacterial communities residing in the rhizosphere confer benefits to their plant host when challenged by biotic and abiotic factors. These communities are modulated by the plant’s root exudates released into the soil. Our study indicated that disruption of the IG pathway caused a shift in the microbial population of the different plant lines. The root exudate profile of *cyp79B2cyp79B3* was significantly different from Col-0 and *atr1D* but there were no differences between the profiles of *atr1D* and Col-0. Nonetheless, both mutant lines displayed a higher relative abundance of beneficial bacterial taxa when compared to Col-0. This suggests that disrupting the IG pathway, whether it is by upregulating it or knocking it out, modulates the presence and abundance of the rhizobacterial community. These findings could have implications for future work focused on improving the relationship between soil microbes and important agricultural crops.

## SUPPLEMENTAL FIGURES

**Figure S1:**
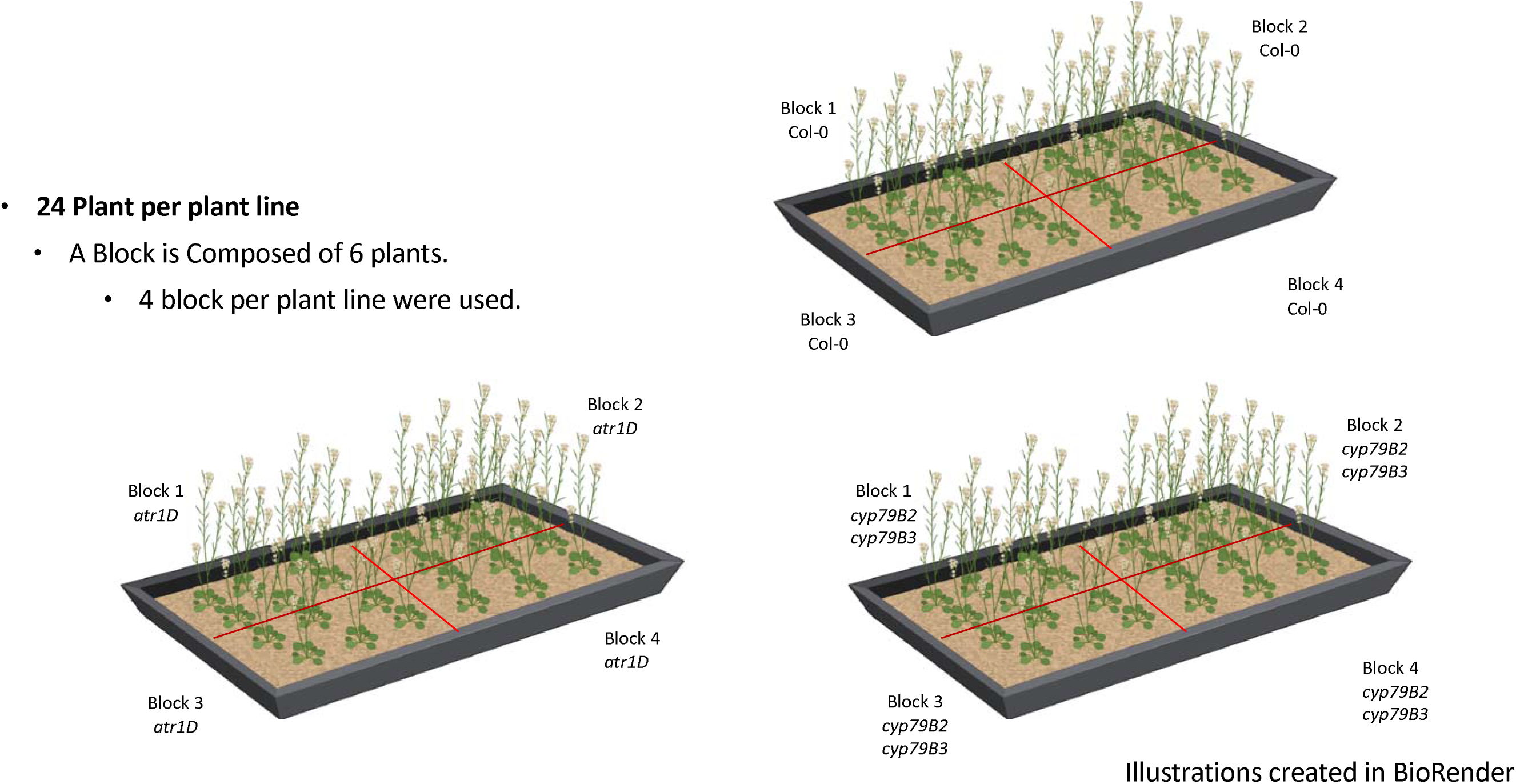
Block design approach.

**Figure S2:**
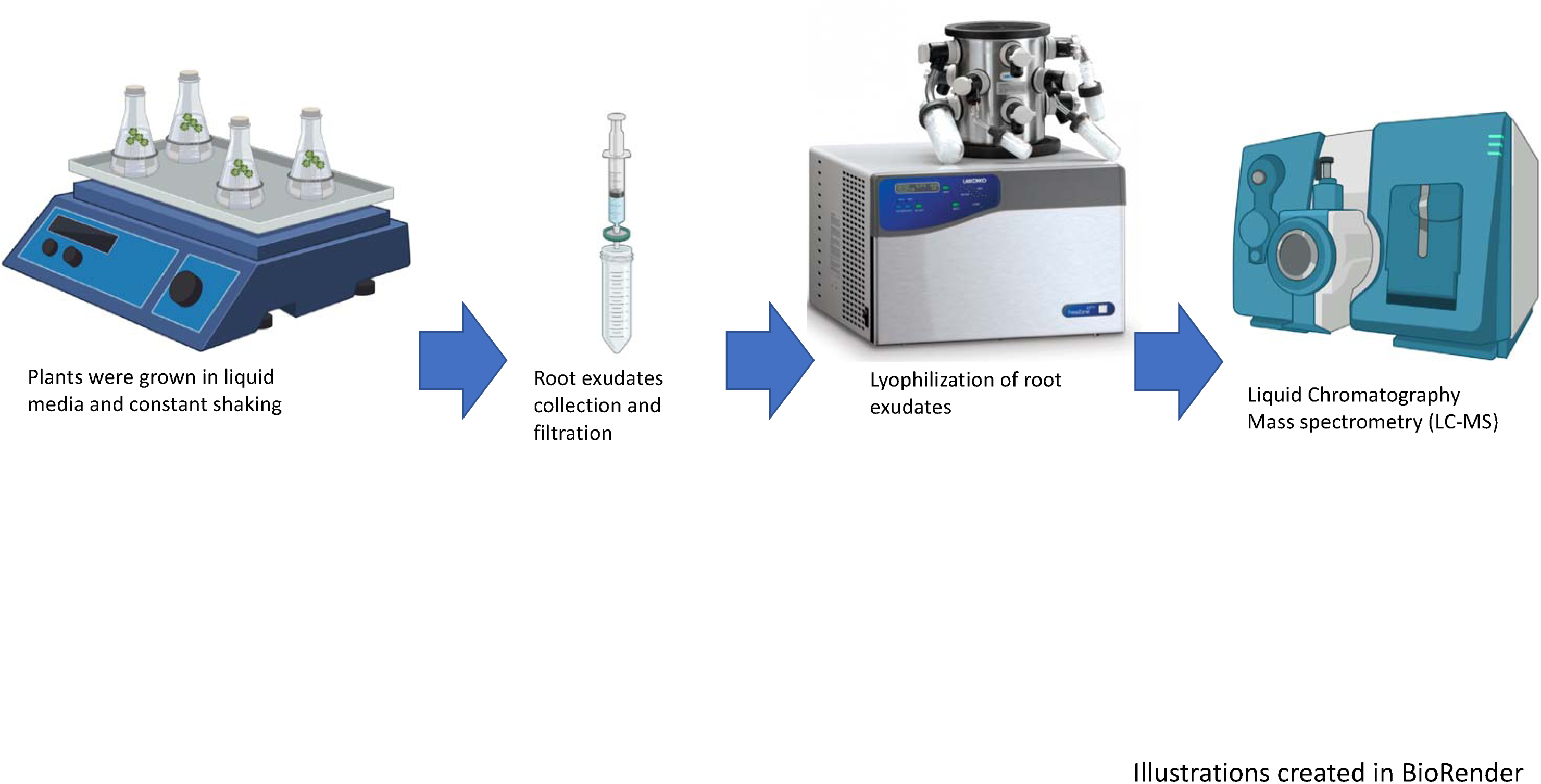
Experimental set up for the collection of root exudates from the different plant lines.

**Figure S3.**
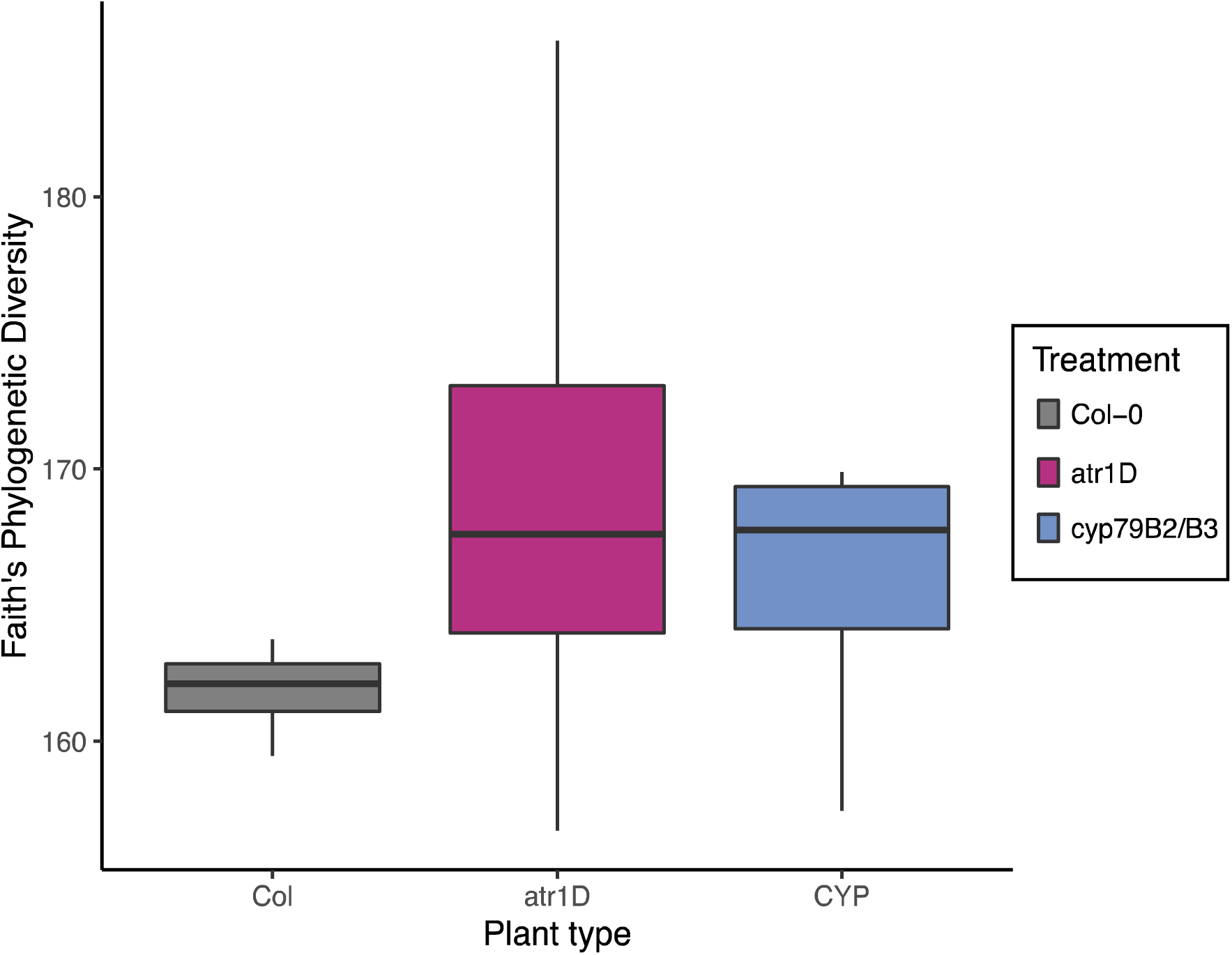
Faith’s phylogenetic diversity analysis showed no significant differences between rhizobacterial communities from the different plant lines (LR Chi-square = 1.89, df = 2, *p*-value = 0.388)

## DECLARATIONS

### AVAILABILITY OF DATA AND MATERIALS

Demultiplexed Illumina sequence data and related metadata have been deposited in the National Center for Biotechnology Information Sequence Read Archive (www.ncbi.nlm.nih.gov/sra) under BioProject IDs: PRJNA1018589.

The raw LC-MS/MS data have been deposited in the Mass Spectrometry Interactive Virtual Environment (MassIVE) database at the Center for Computational Mass Spectrometry, University of California, San Diego under dataset number (MassIVE 000092068, doi:10.25345/C56M33D7Q). The Feature Based Molecular Networking results are available at these GNPS task identifiers.

For the negative mode:

https://gnps.ucsd.edu/ProteoSAFe/status.jsp?task=2a089ce9f8e84738878233cf0090470e For the positive mode:
https://gnps.ucsd.edu/ProteoSAFe/status.jsp?task=bac8fe7bf2f748e49cecc891cce06110 The source code of the Metatlas toolbox used to obtain the extracted ion chromatograms and peak height in this study is openly available from the GitHub repository accessible at https://github.com/biorack/metatlas.

### AUTHORS’ CONTRIBUTIONS

**DA, SAM** and **ACC:** Conceptualization**; DA:** experimental design, data collection and data curation, initial analysis, methodology, visualization, and writing, reviewing, and editing manuscript**; MCB:** formal analysis, visualization, reviewing and editing manuscript**; JS:** data collection and curation, methodology, resources, and writing, reviewing, and editing manuscript**; SK** and **BPB:** formal analysis, visualization, reviewing and editing manuscript**; ACC and TRN:** resources, funding, and writing, reviewing and editing manuscript. All authors read and approved the final manuscript.

## ACKNOWLEDGMENTS

We thank all contributors involved in the completion of this work for their constant support and input, especially Dr. Bletz and Dr. Sasse. We thank the Woodhams lab at the University of

Massachusetts, Boston, and Dr. LaBumbard for sharing their equipment and facilities.

